# Identification and analysis of genomic regions influencing leaf morpho-physiological traits related to stress responses in *Dioscorea alata*

**DOI:** 10.1101/2023.10.18.562915

**Authors:** Komivi Dossa, Mahugnon Ezékiel Houngbo, Jean-Luc Irep, Hâna Chair, Denis Cornet

## Abstract

**Background:** Yams (*Dioscorea* spp.) are significant food security crops especially in West Africa. With the increasing tuber demand and climate change challenges, it is pertinent to strengthen breeding programs for developing high-yielding cultivars with climate resilience. The current study aimed at deciphering the genetic basis of leaf traits related to stress responses in a diverse panel of *Dioscorea alata* genotypes.

**Results:** Phenotypic characterization of 12 traits, including leaf dry matter content, leaf area, net photosynthesis, transpiration rate, transpiration use efficiency, stomatal density, stomatal index, node number, leaf thickness, competitor, stress-tolerator, ruderal (*CSR*) ecological strategy spectrum emphasized significant variations among the genotypes and across two planting locations. Weak correlations were observed among most of traits, suggesting that breeding simultaneously for some of these stress response-related traits would be possible. Heritability was highest for transpiration rate, leaf area and stomatal density, while it was lowest for stress-tolerator, ruderal ecological strategies. Genome-wide association study (GWAS) using high-quality single nucleotide polymorphism (SNPs) identified 24 significant associations on 11 chromosomes, where the association signals were consistent across two locations for traits with high heritability, viz., stomatal density (Chr18) and transpiration rate (Chr3). Further characterization of the significant signals and their related alleles identified advantageous alleles contributing positively to the studied traits. Moreover, 44 putative candidate genes were identified. *Dioal.18G049300* (3 *keto acyl-coenzyme A synthase)* was identified as a strong candidate gene for stomatal density, while *Dioal.12G033600* (*Phosphatidyl inositol monophosphate 5 kinase 4*) was identified for net photosynthesis.

**Conclusion:** Taken together, GWAS and allele segregation analysis for key SNPs provided significant insights into the marker-trait associations, which can be further utilized in breeding programs to improve climate resilience in greater yam.

## Background

Crop improvement involves population development, identification of key traits, phenotyping, deciphering the genetics, and combining the traits of interest [1-3]. Although classical breeding is still an effective way for crop improvement, modern technologies such as marker-trait associations have fast-tracked the understanding of natural variation pertaining phenotypic and genotypic diversities for further utilization in breeding programs [4-7]. Screening phenotypes with genotypic differences that can be manipulated is of fundamental interest in crop improvement. The inherited relationship between phenotypic and genotypic differences has been extensively studied in plants to understand the complex genetic architecture underlying agronomically important traits [8-10]. Phenotypic variation can be traced back to causative loci using quantitative trait loci (QTLs) and association mapping approaches [11]. Genome-wide association studies (GWAS) enable us to overcome some limitations of the QTL mapping approach and provide a relatively high resolution to comprehend allelic diversity [12]. GWAS have been extensively employed to identify the causative variations underlying specific traits in many crops, such as wheat [13, 14], rice [15, 16], cotton [17, 18], maize [19, 20], potato [21, 22] and yams [23-26]. For instance, Gatarira et al. [27] identified several putative candidate genes associated with dry matter accumulation and oxidative browning in *Dioscorea alata* using GWAS. Cormier et al. [28] conducted GWAS to decipher the genetic regulators of flowering control and sex determination in greater yam. Recently, Dossa et al. [29] located the genomic regions controlling tuber flesh color and oxidative browning in *D. alata*.

Yams (*Dioscorea* spp.) are considered major food security crops in Africa and other regions in the world [29, 30]. *Dioscorea rotoundata*, *D. cayenensis*, and *D. alata* are the most important yam species in West and Central Africa [31]. *Dioscorea alata* L., commonly known as greater yam or water yam, is widely distributed worldwide and significantly contributes to food security [27, 32]. In West Africa, greater yam is mainly cultivated in three ecological zones, including rainforests, the southern Guinea savanna, and the wetter portion of the northern Guinea savanna [33]. Rainfalls are the primary source of irrigation in yam-cultivated areas in Africa and 1000 to 1500 mm rainfall during the cropping season favors its production [33]. Leaf development, tuber initiation, and bulking are the critical growth stages, and a lack of sufficient water can reduce the yield to critical levels [34]. Diby et al. [35] reported a significant yield decrease in *D. alata* under water stress conditions. Moisture stress can directly impact dry matter accumulation due to low photosynthetic rate and metabolism, indirectly impacting tuber initiation and development [36, 37]. There is a consensus on developing drought and heat-tolerant greater yam cultivars for stable and improved tuber yield performance [36]. Climate change, unpredicted rainfalls, and increasing temperatures constantly threaten yam production [38, 39]. Moreover, the lack of studies concerning abiotic stress tolerance in yam requires attention to cope with the emerging challenges associated with climate change and food security [37].

Stress tolerance is a multi-stage phenomenon comprising stress sensing, signaling, and recovery [40]. Plants tend to avoid water stress by reducing water demand through avoidance mechanisms that include but are not limited to stomatal closure, plant architecture adjustments, reduced leaf growth, and early leaf senescence [41]. Drought reduces leaf growth [42], rate of photosynthesis [43], stomatal density [44], and yield [45]. Several traits including root structure, leaf morphology, photosynthetic response, osmotic adjustment, water potential, cell membrane stability, and enzymatic activity contribute to stress tolerance [46-48]. Water stress can significantly impact plant growth by hampering morpho-physiological attributes such as photosynthesis, nutrient transport, and carbon assimilation [49, 50]. Leaves are the most sensitive indicators of environmental changes causing abiotic stress [51]. Stomata, as a medium of gaseous exchange, play a crucial role in maintaining immunity and development through the carbon and water cycle in plants [52, 53]. Growth inhibition due to low photosynthesis under water stress is mainly caused by stomatal or non-stomatal attributes such as decreased CO_2_ and low photosynthetic activity in mesophyll tissues [54, 55]. Previous studies have emphasized that leaf traits such as leaf size, thickness, photosynthesis, respiration rate, and stomatal density play an important role in plants exposed to abiotic stress [56-59]. For instance, Li et al. [60] demonstrated that spatiotemporal regulation of *DUF538*, *TRA2,* and *AbFH2* play pivotal roles in leaf physiology in response to water stress. Karaba et al. [61] functionally characterized *HARDY* which improves drought stress tolerance by increasing the photosynthesis and water use efficiency in *Arabidopsis thaliana*.

Research to unravel the genetic basis of major agronomic traits associated with stress response will have far-reaching implications for the genetic improvement of *D. alata*. This is the first such study aimed at deciphering the genetic architectures of leaf morpho-physiological attributes related with stress response in *D. alata*. To achieve this objective, a diversified panel of *D. alata* was assembled and phenotyped for the studied traits at two locations. GWAS was employed to detect the genomic variations and putative candidate genes associated with the studied traits. The results provide a framework for further improvement in breeding programs concerning the development of climate-resilient *D. alata* cultivars.

## Results

### Leaf morpho-physiological characterization

Trait characterization and underlying variations are critical for marker-trait association studies. Hereby, we assembled an association panel composed of 53 genotypes of *Dioscorea alata* as described by Dossa et al. [29] and characterized 12 leaf morpho-physiological traits related to stress responses at two planting sites in Guadeloupe (Duclos and Godet), including leaf dry matter content (LDMC), leaf area (LA1), net photosynthesis (A), transpiration rate (E), transpiration use efficiency (TUE), stomatal density (DS), stomatal index (IS), node number (NN), leaf thickness (LT), competitor, stress-tolerator, ruderal (*CSR*) ecological strategies. Summary statistics showed significant variation for these traits in the association panel, which is useful for deciphering their genetic architectures (Table 1 and Figure S1). The broad-sense heritability (H^2^) computed for the different traits ranged from 0.48 (S) to 0.94 (E).

**Table 1.**
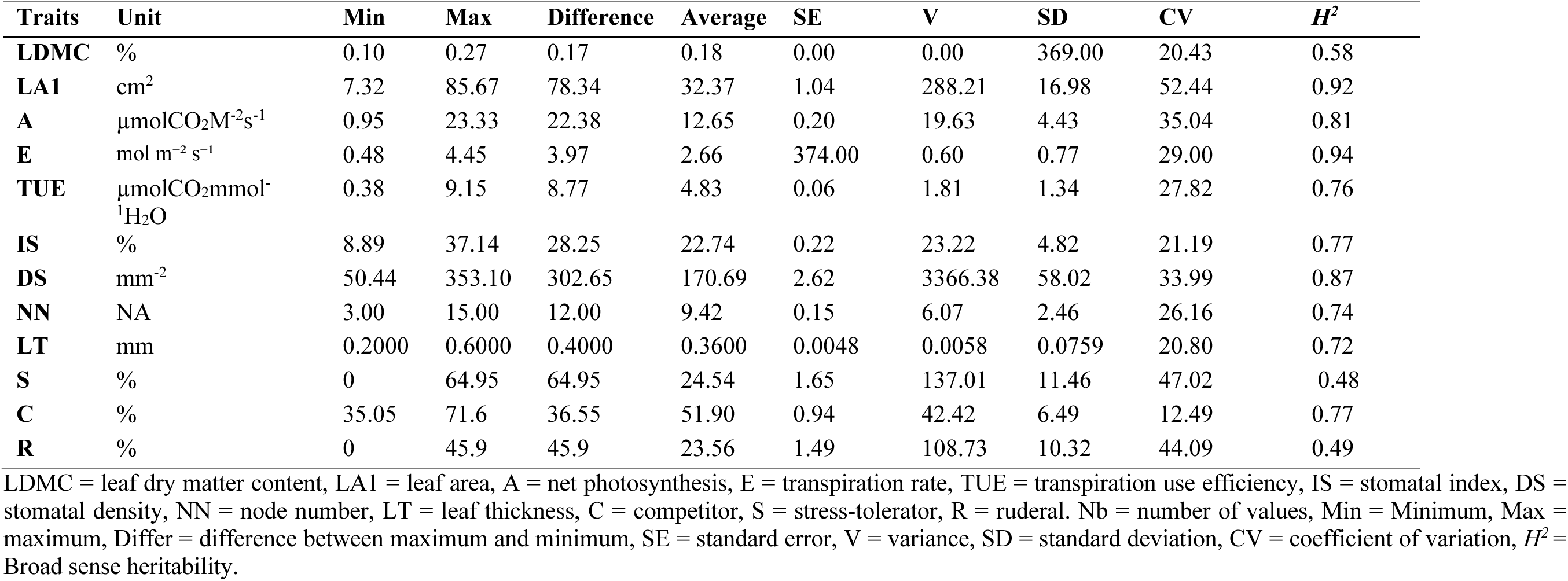
Summary statistics for leaf morpho-physiological traits associated with stress response in greater yam.

| Traits | Unit | Min | Max | Difference | Average | SE | V | SD | CV | $H^2$ |
| --- | --- | --- | --- | --- | --- | --- | --- | --- | --- | --- |
| **LDMC** | % | 0.10 | 0.27 | 0.17 | 0.18 | 0.00 | 0.00 | 369.00 | 20.43 | 0.58 |
| **LA1** | cm<sup>2</sup> | 7.32 | 85.67 | 78.34 | 32.37 | 1.04 | 288.21 | 16.98 | 52.44 | 0.92 |
| **A** | μmolCO<sub>2</sub>M<sup>-2</sup>s<sup>-1</sup> | 0.95 | 23.33 | 22.38 | 12.65 | 0.20 | 19.63 | 4.43 | 35.04 | 0.81 |
| **E** | mol m<sup>-2</sup> s<sup>-1</sup> | 0.48 | 4.45 | 3.97 | 2.66 | 374.00 | 0.60 | 0.77 | 29.00 | 0.94 |
| **TUE** | μmolCO<sub>2</sub>mmol<sup>-1</sup>H<sub>2</sub>O | 0.38 | 9.15 | 8.77 | 4.83 | 0.06 | 1.81 | 1.34 | 27.82 | 0.76 |
| **IS** | % | 8.89 | 37.14 | 28.25 | 22.74 | 0.22 | 23.22 | 4.82 | 21.19 | 0.77 |
| **DS** | mm<sup>-2</sup> | 50.44 | 353.10 | 302.65 | 170.69 | 2.62 | 3366.38 | 58.02 | 33.99 | 0.87 |
| **NN** | NA | 3.00 | 15.00 | 12.00 | 9.42 | 0.15 | 6.07 | 2.46 | 26.16 | 0.74 |
| **LT** | mm | 0.2000 | 0.6000 | 0.4000 | 0.3600 | 0.0048 | 0.0058 | 0.0759 | 20.80 | 0.72 |
| **S** | % | 0 | 64.95 | 64.95 | 24.54 | 1.65 | 137.01 | 11.46 | 47.02 | 0.48 |
| **C** | % | 35.05 | 71.6 | 36.55 | 51.90 | 0.94 | 42.42 | 6.49 | 12.49 | 0.77 |
| **R** | % | 0 | 45.9 | 45.9 | 23.56 | 1.49 | 108.73 | 10.32 | 44.09 | 0.49 |

LDMC = leaf dry matter content, LA1 = leaf area, A = net photosynthesis, E = transpiration rate, TUE = transpiration use efficiency, IS = stomatal index, DS = stomatal density, NN = node number, LT = leaf thickness, C = competitor, S = stress-tolerator, R = ruderal. Nb = number of values, Min = Minimum, Max = maximum, Differ = difference between maximum and minimum, SE = standard error, V = variance, SD = standard deviation, CV = coefficient of variation, $H^2$ = Broad sense heritability.

Furthermore, comparisons of these traits at the two planting locations suggested the planting sites significantly affected TUE, E, LT, S, and R variations within the panel (Figure 1). Pearson’s correlation between traits was estimated, where negative and strong correlations were observed between E and TUE; S and R (Figure 2A). LA1 was positively correlated with C, while negatively correlated with R (Figure 2A). Interestingly, we observed weak correlations between most of traits, suggesting that breeding simultaneously for some of these stress responses-related traits would be possible.

**Figure 1.**
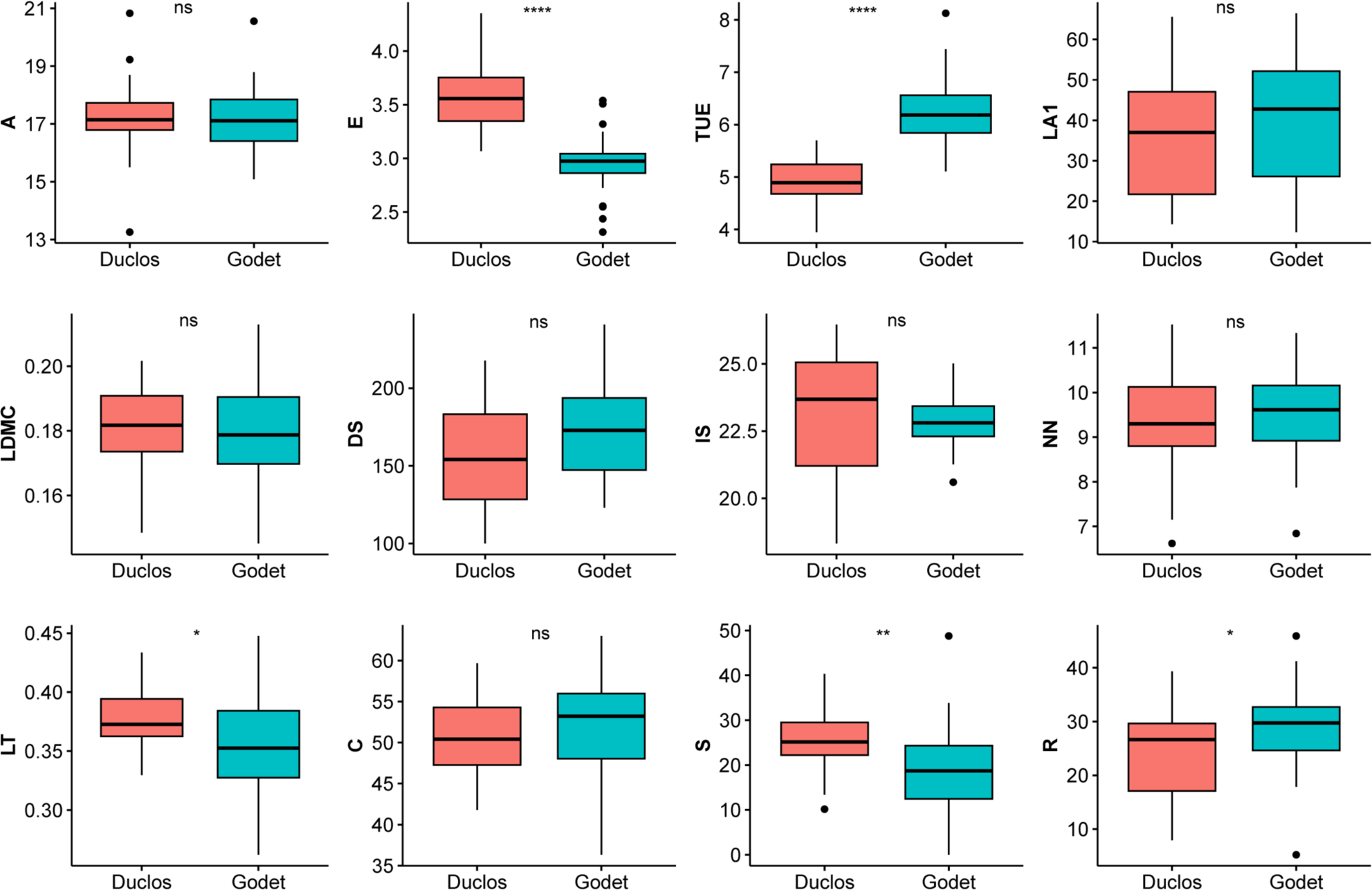
Leaf morpho-physiological trait characterization at two planting sites, Duclos and Godet. Where LDMC = leaf dry matter content, LA1 = leaf area, A = net photosynthesis, E = transpiration rate, TUE = transpiration use efficiency, IS = stomatal index, DS = stomatal density, NN = node number, LT = leaf thickness, C = competitor, S = stress-tolerator, R = ruderal ecological strategies. *Significant difference at *P*=0.05, ****significant difference at *P*=0.001, and ns refers to statistically non-significant difference.

**Figure 2.**
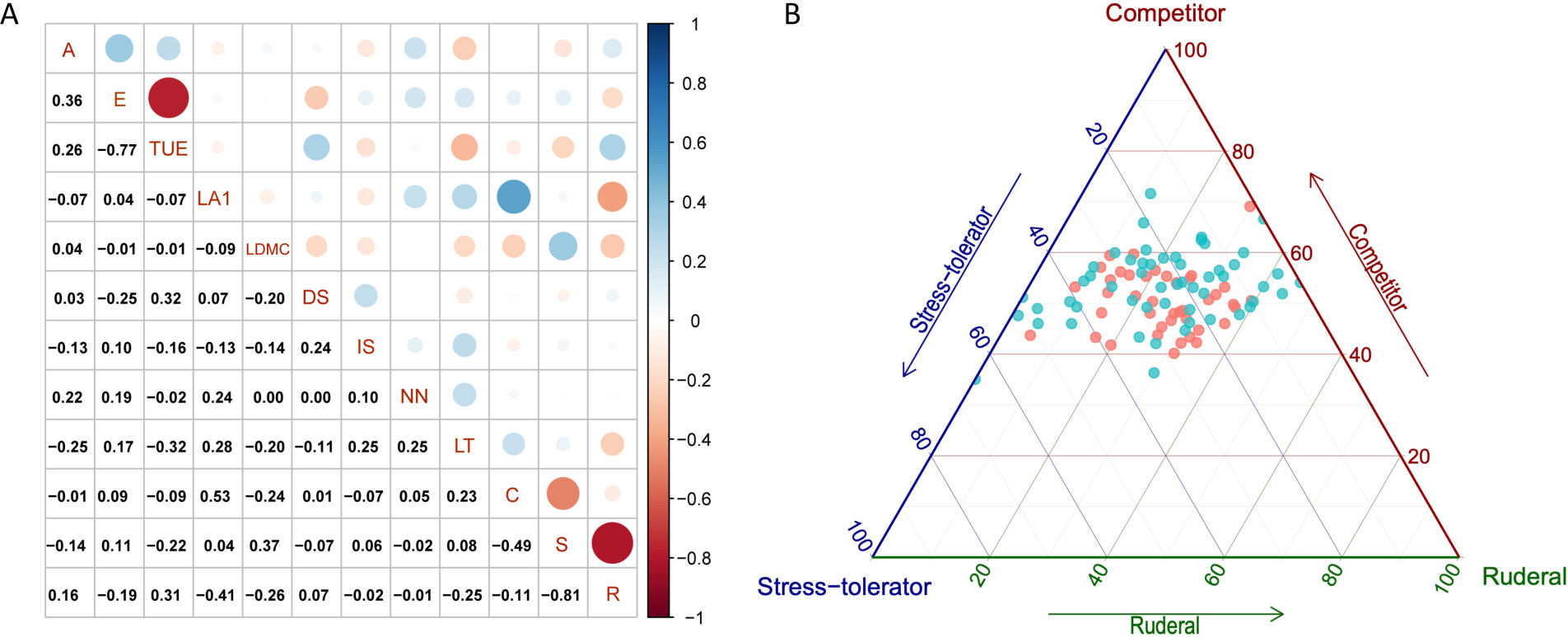
Correlation among traits and overview of the competitor, stress-tolerator, ruderal (*CSR*) ecological strategies in our diversity panel. A) Pearson’s correlation estimates among studied traits; B) Position of yam genotypes of the *D. alata* diversity panel on Grime’s CSR triangle. Orange dots correspond to dryer site (Godet) and purple dots from wetter site (Duclos); Where LDMC = leaf dry matter content, LA1 = leaf area, A = net photosynthesis, E = transpiration rate, TUE = transpiration use efficiency, IS = stomatal index, DS = stomatal density, NN = node number, LT = leaf thickness, C = competitor, S = stress-tolerator, R = ruderal ecological strategies.

#### Erreur ! Source du renvoi introuvable

B represents the ecological strategy of yam genotypes as classified by Grime into competitive ability (C), physiological tolerance to stress (S) and ruderality, i.e., adaptation to disturbance (R). There is quite a large diversity of physiological tolerance to stress (S-R axes) and the genotypes exhibited an important phenotypic plasticity illustrated by different behaviors in the two environments (Figure 2B).

### Genotyping and genome-wide association studies

Whole genome sequencing was utilized for genotyping, and the clean reads were mapped to the reference genome of *D. alata* v2.1 [62]. A total of. 1.9 million high-quality SNPs were used for association studies using the advanced multiple loci model statistical model Bayesian-information and linkage-disequilibrium iteratively nested keyway (BLINK) implemented in the R package GAPIT3 [63]. It has been proven more robust than the traditional Mixed Linear Model [64]. In this study, the genome-wide association study (GWAS) was independently conducted with phenotypic data from each location. The threshold for significant associations was set at log_10_ *P* = 6.

At the two locations, we identified significant associations for all the traits, except for R. We observed that for traits with high *H^2^*, similar genomic regions were stably identified at the two locations. For example, for DS, which has an *H^2^* = 0.87, a strong SNP rs1744535 at position 22836023 on Chromosome (Chr) 18 was identified at Godet (Figure 3A and Table 2) while a close SNP rs1737071 at position 22259955 on Chr18 was detected at Duclos (Figure 3B). Likewise, for E, which has a very high *H^2^*= 0.94 in our panel, GWAS resulted in a significant SNP rs205735 at position 326438 on the Chr3 at Duclos and a close SNP rs202363 at position 346038 on Chr3 at Godet (Figure 3E-3H).

**Figure 3.**
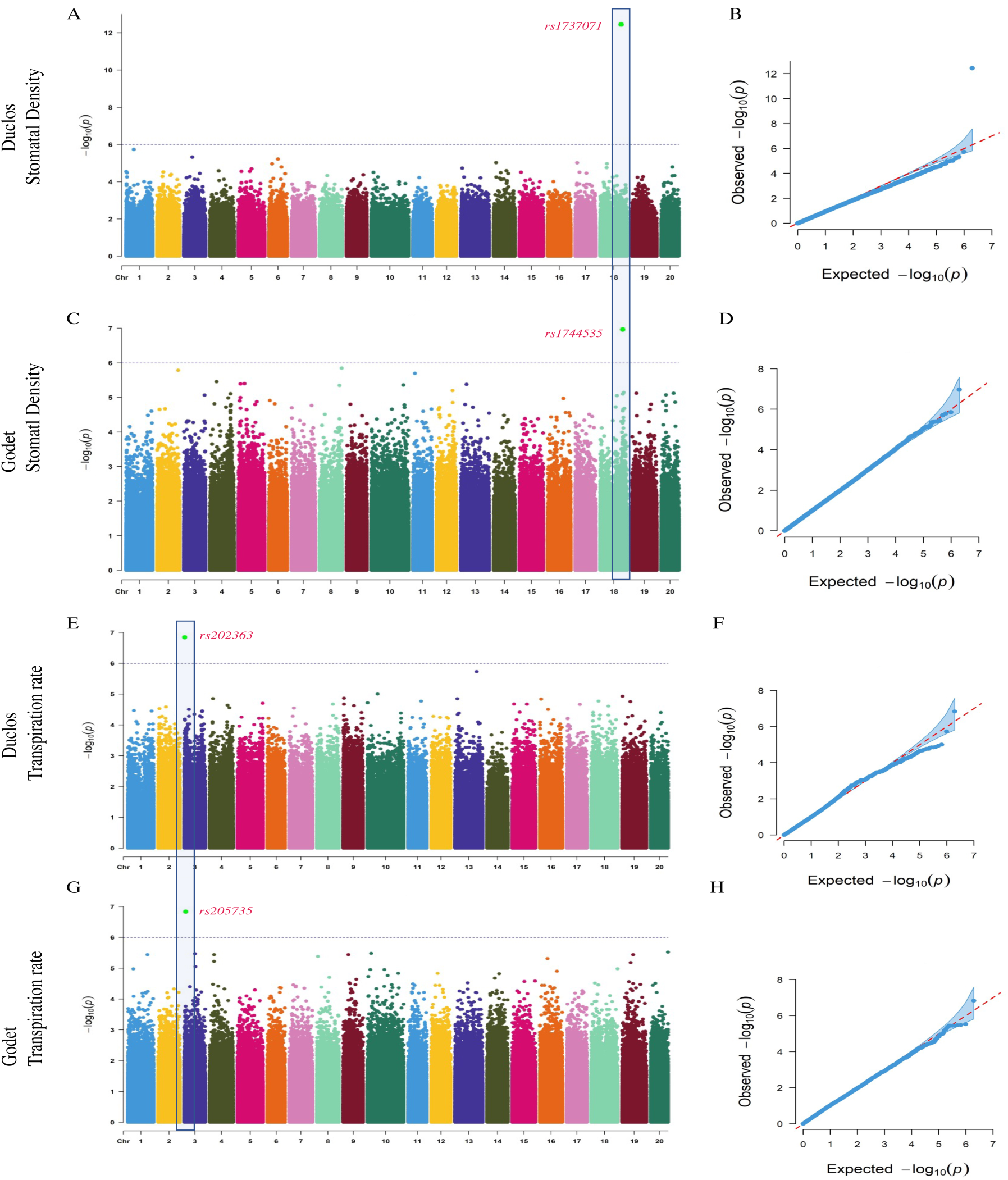
GWAS for stomatal density (DS) and transpiration rate (E) in *D. alata*. A) Manhattan plot for DS at Duclos, with the peaks indicating significant GWAS signals and the dotted horizontal lines indicating the genome-wide significance threshold, B) The QQ-Plot associated with DS at Duclos shows the -log_10_*P* of the expected vs. observed P values of each SNP (blue dots). The red line is a guide for the perfect fit to -log_10_*P*. The shaded area shows the 95% confidence interval for the QQ-plot under the null hypothesis of no association between the SNP and the trait, C) Manhattan plot for DS at Godet, D) The QQ-plot associated with the DS at Godet. E) Manhattan plot for transpiration rate at Duclos, with the peaks indicating significant GWAS signals, and the dotted horizontal lines indicating the genome-wide significance threshold, F) The QQ-Plot associated with transpiration rate at Duclos shows the -log_10_*P* of the expected vs. observed P values of each SNP (blue dots). G) Manhattan plot for transpiration rate at Godet, H) The QQ-plot associated with the transpiration rate at Godet.

**Table 2.** Significant SNPs and their effect on stress response-related leaf traits in *D. alata*.

| Location | Traits | SNP | Chr | Position | Allele | p-value ($-\log_{10}$) | Effect (R<sup>2</sup>) |
| --- | --- | --- | --- | --- | --- | --- | --- |
| **Godet** | LMDC | rs1185429 | Chr13 | 7719492 | G/A | 6.18E+00 | 9.39% |
| **Godet** | IS | rs264342 | Chr3 | 12192098 | T/C | 6.07E+00 | 15.89% |
| **Godet** | E | rs205735 | Chr3 | 1346038 | T/C | 6.83E+00 | 25.28% |
| **Godet** | DS | rs1744535 | Chr18 | 22836023 | A/G | 6.96E+00 | 38.36% |
| **Godet** | A | rs1298027 | Chr14 | 6889304 | C/T | 6.10E+00 | 23.84% |
| **Godet** | C | rs35673 | Chr1 | 9542330 | **G/A** | 7.47E+00 | 31.44% |
|  |  | rs1938821 | Chr20 | 11950212 | T/C | 6.52E+00 | 10.82% |
|  |  | rs1570714 | Chr17 | 1304546 | A/G | 6.34E+00 | 10.49% |
|  |  | rs214171 | Chr3 | 2944668 | **C/T** | 6.00E+00 | 12.85% |
| **Duclos** | TUE | rs304565 | Chr3 | 19525040 | A/G | 6.14E+00 | 12.11% |
|  |  | rs185442 | Chr2 | 18993473 | **G/A** | 6.11E+00 | 19.56% |
|  |  | rs1894629 | Chr20 | 1613539 | G/A | 6.06E+00 | 9.60% |
| **Duclos** | NN | rs1587198 | Chr17 | 4889485 | G/A | 6.40E+00 | 19.89% |
|  |  | rs1254341 | Chr13 | 24382368 | T/C | 6.26E+00 | 16.32% |
|  |  | rs1414555 | Chr15 | 13757209 | T/C | 6.12E+00 | 14.68% |
|  |  | rs1330927 | Chr14 | 14848380 | A/T | 6.07E+00 | 10.05% |
| **Duclos** | LT | rs1091838 | Chr12 | 5760157 | G/A | 1.21E+01 | 31.32% |
| **Duclos** | LDMC | rs450349 | Chr5 | 9839496 | G/A | 6.25E+00 | 11.36% |
| **Duclos** | LA1 | rs401519 | Chr5 | 80613 | **C/T** | 7.19E+00 | 22.84% |
| **Duclos** | IS | rs1436584 | Chr15 | 19347003 | T/A | 6.20E+00 | 15.80% |
| **Duclos** | E | rs202363 | Chr3 | 326438 | **G/A** | 6.84E+00 | 9.23% |
| **Duclos** | DS | rs1737071 | Chr18 | 21259955 | G/A | 1.24E+01 | 36.09% |
| **Duclos** | S | rs1610426 | Chr17 | 9692719 | A/T | 6.28E+00 | 22.19% |
|  |  | rs1111390 | Chr12 | 10073121 | T/G | 6.13E+00 | 24.42% |

LDMC = leaf dry matter content, LA1 = leaf area, A = net photosynthesis, E = transpiration rate, TUE = transpiration use efficiency, IS = stomatal index, DS = stomatal density, NN = node number, LT = leaf thickness, C = competitor, S = stress-tolerator ecological strategies. The alleles in bold letters correspond to the favorable alleles.

However, this is not true for traits with lower *H^2^*. For instance, for LDMC we identified significant SNPs from different genomic regions (Chr5 and Chr13) at Godet and Duclos (Figure 4A-4D). Similarly, two SNPs on chromosome 15 (rs1436584) and 3 (rs264342) were identified for SI at Godet and Duclos, respectively (Figure 4E-4G). Overall, 24 significant marker-trait associations were detected in this study (Figure S2 and Table 2).

**Figure 4.**
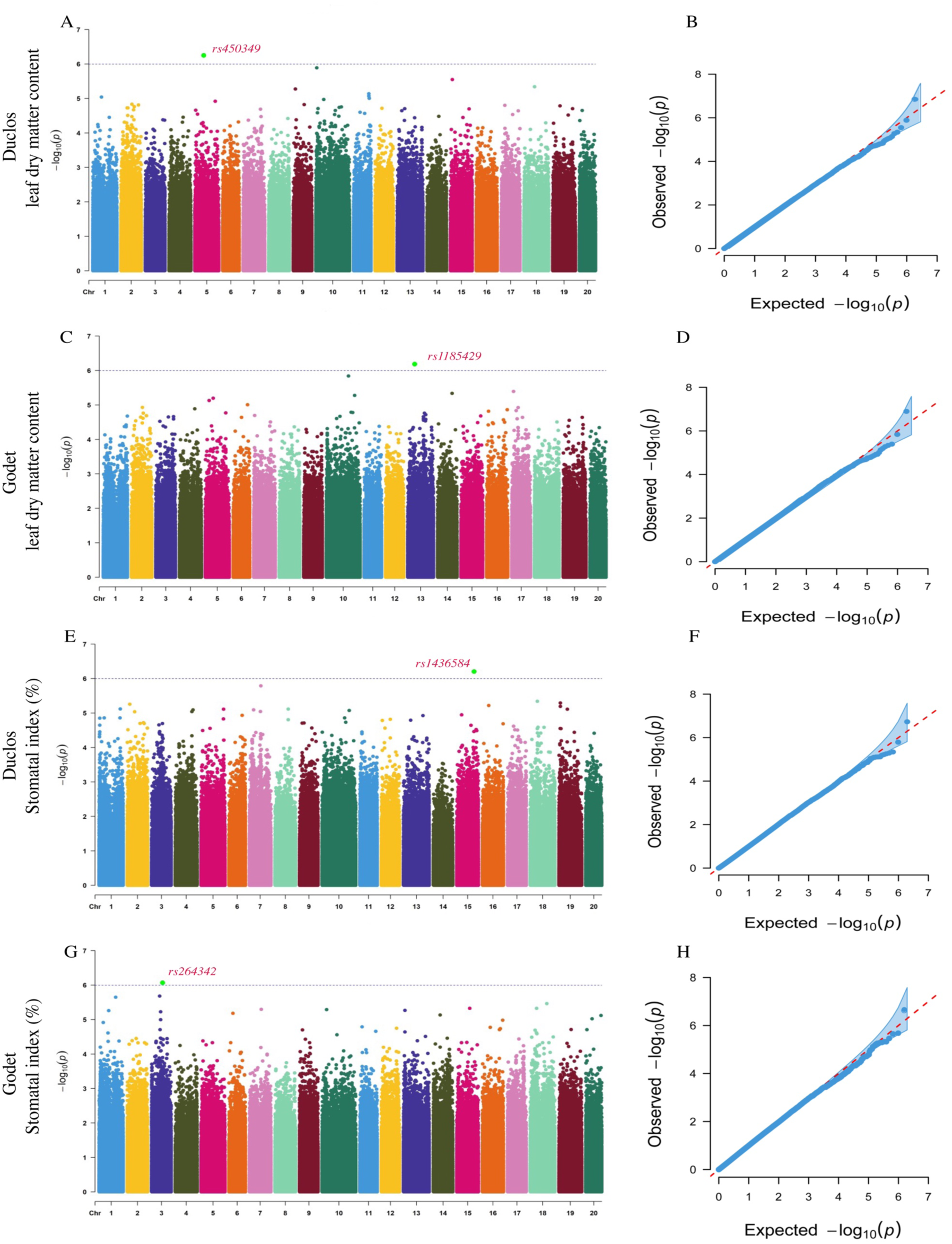
GWAS for leaf dry matter content (LDMC) and stomatal index (IS) in *D. alata*. A) Manhattan plot for LDMC at Duclos, with the peaks indicating significant GWAS signals, and the dotted horizontal lines indicating the genome-wide significance threshold, B) The QQ-Plot associated with LDMC at Duclos shows the -log_10_*P* of the expected vs. observed P values of each SNP (blue dots). The red line is a guide for the perfect fit to -log_10_*P*. The shaded area shows the 95% confidence interval for the QQ-plot under the null hypothesis of no association between the SNP and the trait, C) Manhattan plot for LDMC at Godet, D) The QQ-plot associated with the LDMC at Godet. E) Manhattan plot for IS at Duclos, with the peaks indicating significant GWAS signals, and the dotted horizontal lines indicating the genome-wide significance threshold, F) The QQ-Plot associated with IS at Duclos shows the -log_10_*P* of the expected vs. observed P values of each SNP (blue dots). G) the Manhattan plot for IS at Godet, and H) The QQ plot associated with the IS at Godet.

### Characterization of peak SNPs for their allelic effects on the traits

The favorable alleles at each significant locus were identified and the phenotypic variance explained was computed (Table 2). For example, the SNP (Chr18: rs1737071) could explain 36% of the DS variation in our panel at Duclos (Table 2). We observed that the heterozygote accessions (GA) at this locus had significantly higher DS than the homozygous accessions (GG) (Figure 5). Hence the accessions with the GA genotype could better control water loss rate and CO_2_ uptake [65]. Similarly, SNP (Chr5: rs450349) explained 11.36% of the variation with the allele GA associated with higher LDMC compared to GG (Figure 5). High DS and LDMC have been linked to improved photosynthetic induction and biomass production in plants [66]. Moreover, other peak SNPs detected at Duclos and the nine peak SNPs detected at Godet were characterized for their favorable alleles (Figure 5 and Figure 6).

**Figure 5.**
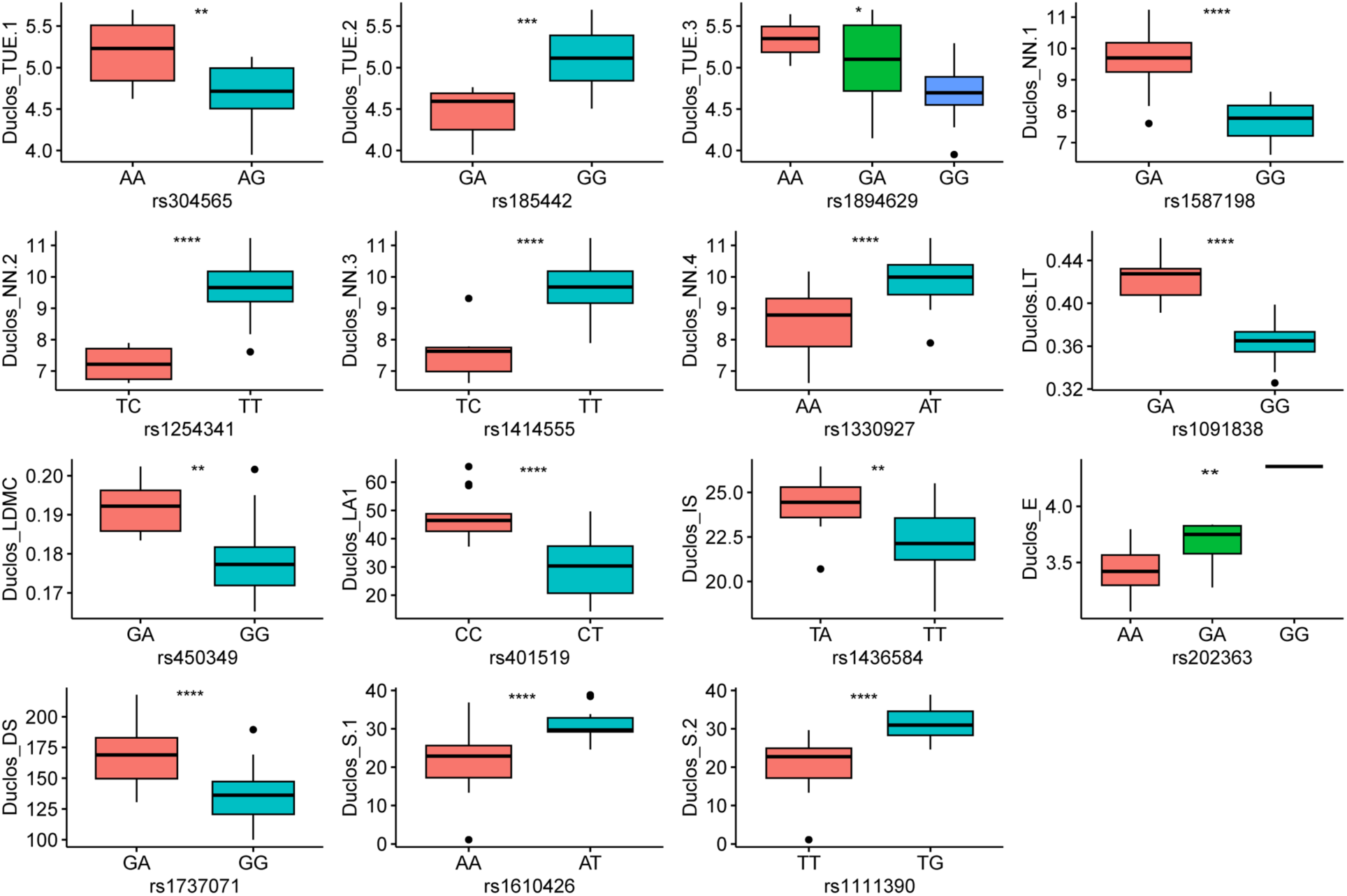
Allelic effect of 15 significant SNPs identified at Duclos. LDMC = leaf dry matter content, LA1 = leaf area, A = net photosynthesis, E = transpiration rate, TUE = transpiration use efficiency, IS = stomatal index, DS = stomatal density, NN = node Number, LT = leaf thickness, S = stress-tolerator ecological strategy. *Significant difference at *P*=0.05, ** significant difference at *P*=0.01, and ****significant difference at *P*=0.001.

**Figure 6.**
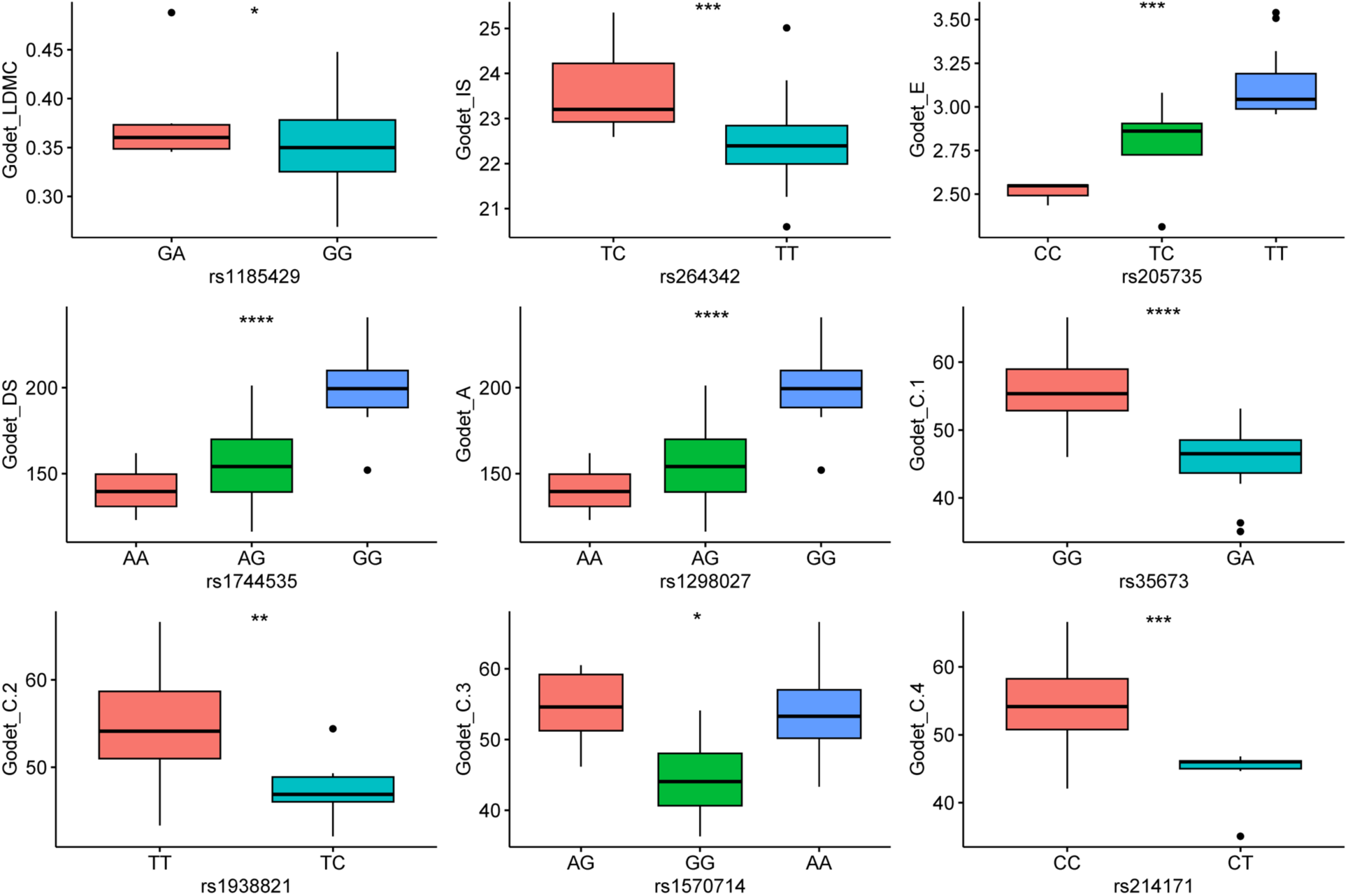
Allelic effect of nine significant SNPs identified at Godet. A = net photosynthesis, DS = stomatal density, E = transpiration rate, IS = stomatal index, LDMC = leaf dry matter content, C = competitor ecological strategy. *Significant difference at *P*=0.05, ** significant difference at *P*=0.01, and **** significant difference at *P*=0.001.

### Identification of candidate genes associated to the stress response leaf traits

We searched for candidate genes controlling the studied traits around the associated significant SNPs. Overall, 44 genes with diverse biological functions were extracted. Sixteen genes with known functions were identified as putative candidates for physiological attributes (DS, A, E, TUE, and IS) associated with stress response. For the DS trait at Duclos, the peak SNP was located between two genes (*Dioal.18G049200* and *Dioal.18G049300*) (Table 3). There was no functional annotation for *Dioal.18G049200,* but the gene *Dioal.18G049300* was annotated as a 3- keto acyl-coenzyme A synthase (*KCS*). At Godet, the peak SNP were identified downstream of *Dioal.18G055600* annotating *GLUTATHIONE S-TRANSFERASE*. We speculate that *Dioal.18G049300* and *Dioal.18G055600* could be the potential regulators of DS in *D. alata*.

**Table 3.** List of candidate genes associated with stress response-related leaf traits in greater yam.

| Traits | SNP | Upstream the gene<br>/downstream the gene | Localization | Function |
| --- | --- | --- | --- | --- |
| Godet_LMDC | rs1185429 | *Dioal.13G049900*<br>*Dioal.13G050000* | intergenic | Retrotransposon protein<br>Neprosin_AP |
| Godet_IS | rs264342 | *Dioal.03G049700*<br>*Dioal.03G049800* | intergenic | SHR-BD<br>Major facilitator superfamily protein |
| Godet_E | rs205735 | *Dioal.03G019500*<br>*Dioal.03G019600* | intergenic | Fatty acid hydroxylase superfamily<br>Nodulin MtN21 /EamA-like transporter family protein |
| Godet_DS | rs1744535 | *Dioal.18G055600* | downstream | GLUTATHIONE S-TRANSFERASE |
| Godet_A | rs1298027 | *Dioal.12G033500*<br>*Dioal.12G033600* | intergenic | ABC-2 type transporter family protein<br>Phosphatidyl inositol monophosphate 5 kinase 4 |
| Godet_C | rs35673 | *Dioal.01G024000*<br>*Dioal.01G024100* | intergenic | IRON/ASCORBATE OXIDOREDUCTASE<br>RNA-directed DNA polymerase |
|  | rs1938821 | *Dioal.20G032000*<br>*Dioal.20G032100* | intergenic | NA<br>PROTEASE DO-LIKE 1 |
|  | rs1570714 | *Dioal.17G018200* | upstream | S-ADENOSYLMETHIONINE CARRIER 2 |
|  | rs214171 | *Dioal.03G033700*<br>*Dioal.03G033800* | intergenic | Leucine Rich Repeat<br>F-box domain and LRR containing protein |
| Duclos_TUE | rs304565 | *Dioal.03G073900*<br>*Dioal.03G074000* | intergenic | Ankyrin repeat family protein<br>Seed imbibition 2 |
|  | rs185442 | *Dioal.02G062100* | exonic | Non-specific serine/threonine protein kinase |
|  | rs1894629 | *Dioal.20G005200*<br>*Dioal.20G005300* | intergenic | Protein phosphatase 2C family protein<br>Non-specific serine/threonine protein kinase |
| Duclos_NN | rs1587198 | *Dioal.17G048200*<br>*Dioal.17G048300* | intergenic | CCT motif family protein<br>UDP-D-glucose/UDP-D-galactose 4-epimerase 5 |
|  | rs1254341 | *Dioal.13G076500*<br>*Dioal.13G076600* | intergenic | Exostosin family<br>NA |
|  | rs1414555 | *Dioal.15G047700*<br>*Dioal.15G047800* | intergenic | Lipid transfer protein 12<br>Non-specific serine/threonine protein kinase |
|  | rs1330927 | *Dioal.14G073400*<br>*Dioal.14G073500* | intergenic | S-adenosyl-L-methionine-dependent methyltransferase protein<br>pentatricopeptide |
| Duclos_LT | rs1091838 | *Dioal.12G033000*<br>*Dioal.12G033100* | intergenic | NA<br>Domain of unknown function (DUF4228) |
| Duclos_LDMC | rs450349 | *Dioal.05G072500*<br>*Dioal.05G072600* | intergenic | Chloroplast sensor kinase<br>NA |
| Duclos_LA1 | rs401519 | *Dioal.05G000800*<br>*Dioal.05G000900* | intergenic | Non-specific serine/threonine protein kinase<br>DNA/RNA helicase protein |
| Duclos_IS | rs1436584 | *Dioal.15G069300*<br>*Dioal.15G069400* | intergenic | Bifunctional inhibitor/lipid-transfer protein/seed storage 2S<br>albumin superfamily protein<br>ARM repeat superfamily protein |
| Duclos_E | rs202363 | *Dioal.03G004300* | intronic | Heavy metal ATPase 4 |
| Duclos_DS | rs1737071 | *Dioal.18G049200*<br>*Dioal.18G049300* | intergenic | NA<br>Very-long-chain 3-ketoacyl-CoA synthase |
| Duclos_S | rs1610426 | *Dioal.17G055400*<br>*Dioal.17G055500* | intergenic | NA<br>Tetratricopeptide repeat domain containing protein |
|  | rs1111390 | *Dioal.12G035200*<br>*Dioal.12G035300* | intergenic | WD domain, G-beta repeat (WD40)<br>Retrotransposon gag protein |

LDMC = leaf dry matter content, LA1 = leaf area, A = net photosynthesis, E = transpiration rate, TUE = transpiration efficiency, IS = stomatal index, DS = stomatal density, NN = node numbers, LT = leaf thickness.

Net photosynthesis showed one association signal rs1298027 on chromosome 14, accounting for 23.84% of the phenotypic variance (Table 2). The peak SNP was located between *Dioal.12G033500* and *Dioal.12G033600,* annotated as *ABC-2 type transporter family protein* and *Phosphatidyl inositol monophosphate 5 kinase 4*, respectively. Moreover, three putative candidate genes (*Dioal.03G019500*, *Dioal.03G019600*, and *Dioal.03G004300*) for E, five (*Dioal.03G073900*, *Dioal.03G074000*, *Dioal.02G062100*, *Dioal.20G005200*, and *Dioal.20G005300*) for TUE, and four (*Dioal.03G049700*, *Dioal.03G049800*, *Dioal.15G069300*, *and Dioal.15G069400*) for IS were identified, some containing exonic and intronic SNPs.

For leaf morphological traits (LDMC, LA1, and LT) and NN, 13 putative candidate genes were identified on different chromosomes. *Retrotransposon protein*, *Neprosin_AP*, and *Chloroplast sensor kinase* were identified as putative candidate genes for LDMC. *Dioal.17G048200, Dioal.17G048300, Dioal.13G076500, Dioal.15G047700, Dioal.15G047800, Dioal.14G073400, and Dioal.14G073500* were identified for NN (Table 3). *Domain of unknown function* (*DUF4228*), *Non-specific serine/threonine protein kinase*, and *DNA/RNA helicase protein* were identified as candidate genes for LA1 and LT.

For C and S traits, we identified 11 genes around the significant SNPs with some genes (IRON/ASCORBATE OXIDOREDUCTASE, F-box domain and S-ADENOSYLMETHIONINE) well known to be involved in stress responses in plants (Table 3).

Gene ontology enrichment analysis of the candidate genes highlighted several terms related to leaf morphogenesis and stress response, including regulation of gene expression, regulation of photosynthesis, wax biosynthetic process, response to water deprivation, cuticle development, stomatal movement, cellular response to starvation, auxin−activated signaling pathway (Figure 7). All these results suggest that the candidate genes detected in this study have potential for enhancing stress response in *D. alata* genotypes.

**Figure 7.**
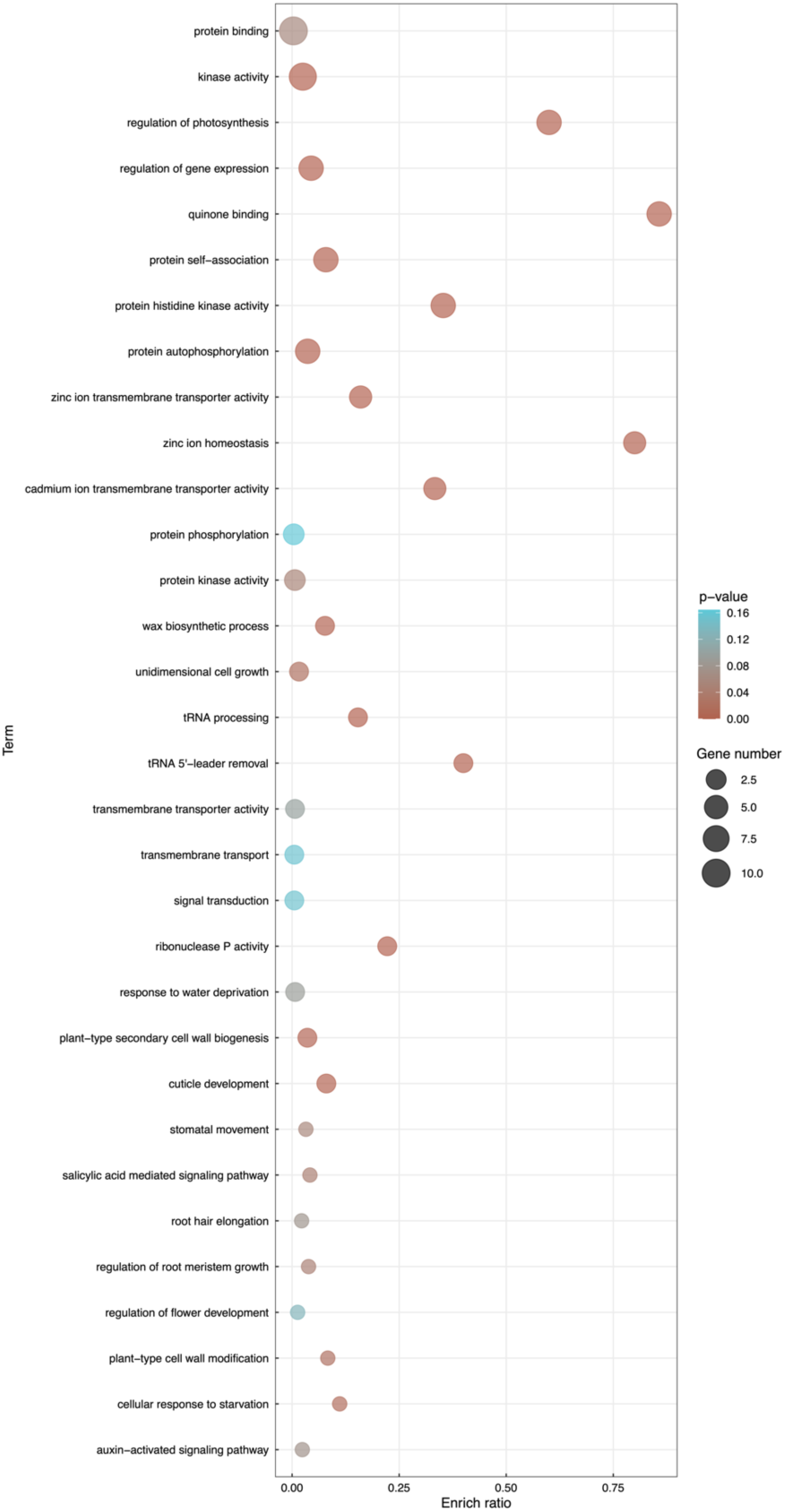
Gene ontology enrichment for the candidate genes detected around the significant SNPs associated with stress response-related leaf traits in greater yam.

## Discussion

In this study, we investigated the genetic basis for several leaf traits related to stress response through genome-wide association studies (GWAS). Although GWAS have been successfully implemented for various traits in yams [23, 25–28, 67, 68], stress-related traits have not yet been characterized using GWAS or other genetic approaches [37].

In this study, GWAS resulted in the identification of 24 significant associations with a stringent threshold (log_10_*P* = 6), which was much higher than previously published reports [23, 25, 28]. A stringent threshold increased the significance of identified SNPs and candidate genes but may have reduced the number of significant associations [7]. Despite successfully identifying genetic variants associated with these traits, the phenotypic variation explained are much lower relative to heritability estimates. This has been reported to be related to the presence of structural variations, genotype-environment interactions, epistasis, origin, and errors in heritability estimates [69, 70]. However, we identified consistent associations at two locations where trait heritability estimates were high (E and DS). These results are consistent with previous reports suggesting the identification of stable QTLs for traits with high heritability [71, 72].

Identification and characterization of alleles associated with key SNPs enable us to understand the effect of significant associations on the leaf traits [7, 73]. Response to abiotic stress is a complex mechanism [74]. During stress, photosynthesis is mainly affected, and plants tend to close their stomata, simultaneously restricting the inflow of CO_2_ and reducing the photosynthesis rate [75]. We identified 24 advantageous alleles with positive contributions to the studied traits. It is important to mention that although the traits studied are known to be involved in stress response in plants, further studies are needed to check how the favorable alleles at the marker-trait associations influence stress response in greater yam genotypes. Hence, abiotic stress experiments can be carried out using selected genotypes in our diversity panel to confirm the relationship between the leaf traits, the alleles at the associated SNPs and stress response. Genotypes with advantageous alleles (associated with trait performance) could be further utilized in breeding programs to accumulate the desirable variation for developing resilient cultivars [76, 77]. Genomic-assisted breeding which has been successful in the development of resilient crops against stress, such as wheat [78, 79] and rice [80, 81] could be applied in greater yam.

The gene *Dioal.18G049300* annotated as 3-keto acyl-coenzyme A synthase (*KCS*) enzyme was identified as a putative candidate gene for DS, which is involved in synthesizing very long-chain fatty acids. It has been demonstrated that the *Arabidopsis thaliana* KCS gene (hic) controls stomatal development through CO_2_ perception, and the mutant plants displayed a 42% increase in stomatal density in response to elevated CO_2_ [82]. Another gene *Dioal.18G055600* was identified, annotated as *GLUTATHIONE S-TRANSFERASE* (*GST*). *GST* has been widely implicated in the response to a wide range of stress conditions, including biotic and abiotic stresses [83-85].

Similarly, *Dioal.12G033500* (*ABC-2 type transporter family protein*) was identified as a putative candidate gene for net photosynthesis. A previous report suggests that disruption of *ATP-binding cassette transporters* causes deregulation of stomatal opening and increases drought susceptibility [86]; therefore, variation in *Dioal.12G033500* function or expression could explain the photosynthesis variation in the GWAS panel and could be an excellent candidate for further functional verification in *D. alata*. *Retrotransposon protein* [87], *Domain of unknown function* (*DUF4228*) [88, 89], *Non-specific serine/threonine protein kinase* [90, 91], *DNA/RNA helicase protein* [92, 93] have also been characterized for their potential role in coping with stress conditions and maintaining the plant growth. Gene expression data facilitate the identification of candidate genes from GWAS projects since genetic variants falling in genic or intergenic regions often lead to non-functional genes or altered gene expression. There is no transcriptome data related abiotic stress response available in greater yam. Therefore, we recommend that a future study can generate gene expression data under various stresses to support the ongoing genetic studies [44].

## Conclusions

We successfully employed GWAS to detect genomic signals (24) and candidate genes (44) linked to leaf traits related to stress response in *Dioscorea alata*. Traits with high heritability estimates, stomatal density, and transpiration rate, were identified with consistent GWAS signals on Chr18 and Chr3. Nonetheless, additional efforts are needed to enlarge the diversity panel to improve the GWAS power and detect more marker-trait associations. Accessions accumulating favorable alleles at the different significant loci will be used as genitors in our breeding programs. Moreover, developing allele-specific markers for the strong and stably detected significant marker-trait associations will improve marker-assisted selection, introgression, and the pyramiding of favorable alleles/genes in greater yam cultivars.

## Methods

### Plant material and leaf morpho-physiological characterization

Plant materials used in this study include 53 genotypes of *Dioscorea alata* L. [28] (Table S1). This panel comprises genotypes from nine countries of the yam belt countries of west Africa, the Caribbean, and the Pacific islands. The plant materials were obtained from the germplasm collection of the Centre de coopération Internationale en Recherche Agronomique pour le Développement, Guadeloupe. The plant materials were planted at two locations, Duclos (16°120′ N, 61°39′ O, 125 m above sea level (a.s.l.)), and Godet (16°20′ N, 61°30′ 0.10 m a.s.l.), in Guadeloupe. A total of 10 seedlings of each cultivar were planted in three replicates (30 seedlings per genotype in total). These plants were spread over three ridges (65 m long) spaced 30 cm apart within the ridges. Cane straw was used as mulch over the entire plot to limit weeds at Godet, while paper mulch was used at Roujol. The plot was drip irrigated, and planting started in March 2021. Genotypes were planted from harvested seeds at year n-1 when 50% of the tubers of the genotype germinated in the storage shed. All the genotypes were characterized for 12 morpho-physiological traits associated with stress response, including leaf dry matter content (LDMC), leaf area (LA1), net photosynthesis (A), transpiration rate (E), transpiration efficiency (TUE), stomatal index (IS), stomatal density (DS), node number from apex to second mature leaf (NN), leaf thickness (LT), C = competitor, S = stress-tolerator, R = ruderal ecological strategies as classified by Grime.

The traits were characterized 17-24 days after the germination when 50% of individuals within each plot had enough leaves for trait measurement. LDMC was measured from three collected leaves for each genotype. Fresh weight and dry weight (wrapped in kraft paper and dried in an oven at 60 °C for three days). LDMC was calculated using the formula.

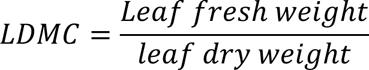

Leaf area was estimated by *Leaf area = length × width*

Vernier caliper was used to estimate LT, and nodes were counted manually to record NN.

### Determination of stomatal density and index

Leaf samples were collected as fresh leaves wrapped in similar-sized wet papers to avoid drying and transported to the laboratory in a cooler. Samples were collected in the morning before 9 am. The stomata are present on the rear side of the leaf [94]. To estimate the stomatal density, the procedure was adapted from Paul et al. [95] to prepare the slides for microscopic observations using suitable replica fluid/adhesive. The slides were then observed under a microscope with 40x magnification, and 60 images were saved for each genotype with ZEN software (Carl Zeiss microscope, GmbH, version 2.3, blue version, 2011). ZEN allows us to directly convert the number of pixels to metric units and get the scale of the captured image. We calculated stomatal density (DS) and stomatal index (IS) by the following formula.

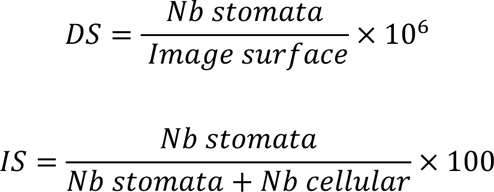

Where *Nb* is the number of stomata per unit area, and *Nb* cellular is the number of epidermal cells per unit area

### Gas exchange measurement

To estimate the gaseous exchange (A, E, TUE), we used ADC LCPro+infrared gas exchange analyzer (ADC bioscience, Hodstone UK Limited). All measurements were taken from 7 am to 9 am in the morning. The parameters were set as follows: temperature 30 °C, saturation brightness is 1400 µ mol photon m^-2^s^-1^, while relative humidity was stable (80%). We estimated A and E and then calculated transpiration use efficiency by the following formula

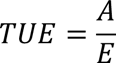

### Genome-wide association studies

Dossa et al. [29] fully detailed the DNA extraction, sequencing and data processing steps. GWAS was performed with all 1.9 million SNPs using the advanced multiple loci model statistical model Bayesian-information and linkage-disequilibrium iteratively nested keyway (BLINK) implemented in the R package “GAPIT3” [63]. BLINK provides an efficient platform with increased power compared to other methods, while reducing the calculation time. The Manhattan plots were also generated in R4.0.23 with the “CMplot” package [99]. SNPs having a significant association with traits were determined by the adjusted p-value. The threshold of *P* < 10^-8^ (0.05/n, with n = number of SNPs) was set to report a significant association. The quantile-quantile (QQ) plots were generated by plotting the negative logarithms (−log_10_) *P*-values relative to their expected p-values to fit model relevance GWAS with the null hypothesis of no association and to determine to what extent the models considered the structure of the population.

### Putative candidate gene identification

To inventory potential genes near associated SNP markers for target traits, we used ANNOVAR version 2.4 to identify genes downstream and upstream of the significant SNPs based on linkage disequilibrium (LD = 5 Kb) [29]. The identified candidate genes were screened for their putative functional attributes. Subsequently, we extracted the related genes and conducted a Gene Ontology (GO) enrichment analysis using the KOBAS-i tool. The significantly enriched GO terms were then visualized using the “ggplot” package from the R4.0.23 software.

### Statistical analysis

The effect of alleles at significant SNPs was assessed by comparing phenotyping data for haplotype groups. A Student’s t-test was used to compare the groups of haplotypes (*P* < 0.05) in the R4.0.23 software with the “ggpubr” and “rstatix” packages.

Using the LME4 package [100] available on R4.0.23, we considered the genotype as a random effect to obtain the variance components of all the traits. We calculated phenotypic variance share of genetic source as broad sense heritability (H²):

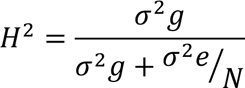

Where *σ*_2_*g* is genotype variance, *σ*_2_*e* is residual variance, and N is the average value of the replicates for each genotype.

The basic descriptive analysis of the stress response-related leaf traits was performed by statistix 8.1. Pearson’s correlation analysis between different stress response-related traits was performed by R4.0.23 using the “corrplot” package [101]. The differences in two planting sites for each trait were estimated using analysis of variance according to Steel et al. [102]. AOV-function in R4.0.23 was employed to perform analysis of variance. *P*-value < 0.05 was regarded as significant.

## Abbreviations

GWAS: Genome wide association studies
SNP: Single nucleotide polymorphism
QTLs: Quantitative trait loci
LDMC: Leaf dry matter content
LA1: Leaf area
A: Net photosynthesis
E: Rate of transpiration
TUE: Transpiration use efficiency
IS: Stomatal index
DS: Stomatal density
NN: Node number
LT: Leaf thickness
C: Competitor ecological strategies
S: Stress-tolerator ecological strategies
R: ruderal ecological strategies
SD: Standard deviation
CV: Coefficient of variation
H^2^: Broad sense heritability
BLINK: Bayesian-information and linkage-disequilibrium iteratively nested keyway
GO: Gene ontology

## Declarations

## Ethics approval and consent to participate

All relevant institutional, national, and international guidelines and legislations were followed while conducting this experiment. The plant materials are available at the germplasm collection of the Centre de coopération Internationale en Recherche Agronomique pour le Développement (CIRAD), Guadeloupe.

## Consent for publication

Not applicable.

## Availability of data and materials

The Illumina NovaSeq 6000 sequencing raw data are available in the NCBI Sequence Read Archive, under the BioProject number: PRJNA880983. The phenotypic datasets are available from the corresponding author upon request.

## Competing interests

The authors declare no conflict of interest.

## Funding

This work was supported by the Bill and Melinda Gates Foundation (BMGF) through the AfricaYam project (grant opportunity INV-003446, formerly OPP1112307).

## Authors’ contributions

*Conceptualization*: Komivi Dossa, Denis Cornet. *Data collection*: Jean-Luc Irep, Denis Cornet *Data curation and Formal analysis*: Komivi Dossa, Mahugnon Ezékiel Houngbo, Denis Cornet. *Funding acquisition*: Denis Cornet, Hâna Chair. *Writing ± original draft*: Komivi Dossa. *Writing ± review & editing*: Denis Cornet, Hâna Chair. All authors have read and approved the final version of this manuscript.

## Acknowledgments

We thank Angélique Morel for helping in data processing. We also thank A. Peter, B. Tessi, J. Bernard, J. Sirejol and Schneyder L. for phenotyping. We also thank C. Perrot and E. Nudol for field assistance and data acquisition.

## Notes

### Competing Interest Statement

The authors have declared no competing interest.

