## Supplementary material for "Identification and analysis of genomic regions influencing leaf morpho-physiological traits related to stress responses in *Dioscorea alata*": Figure S2

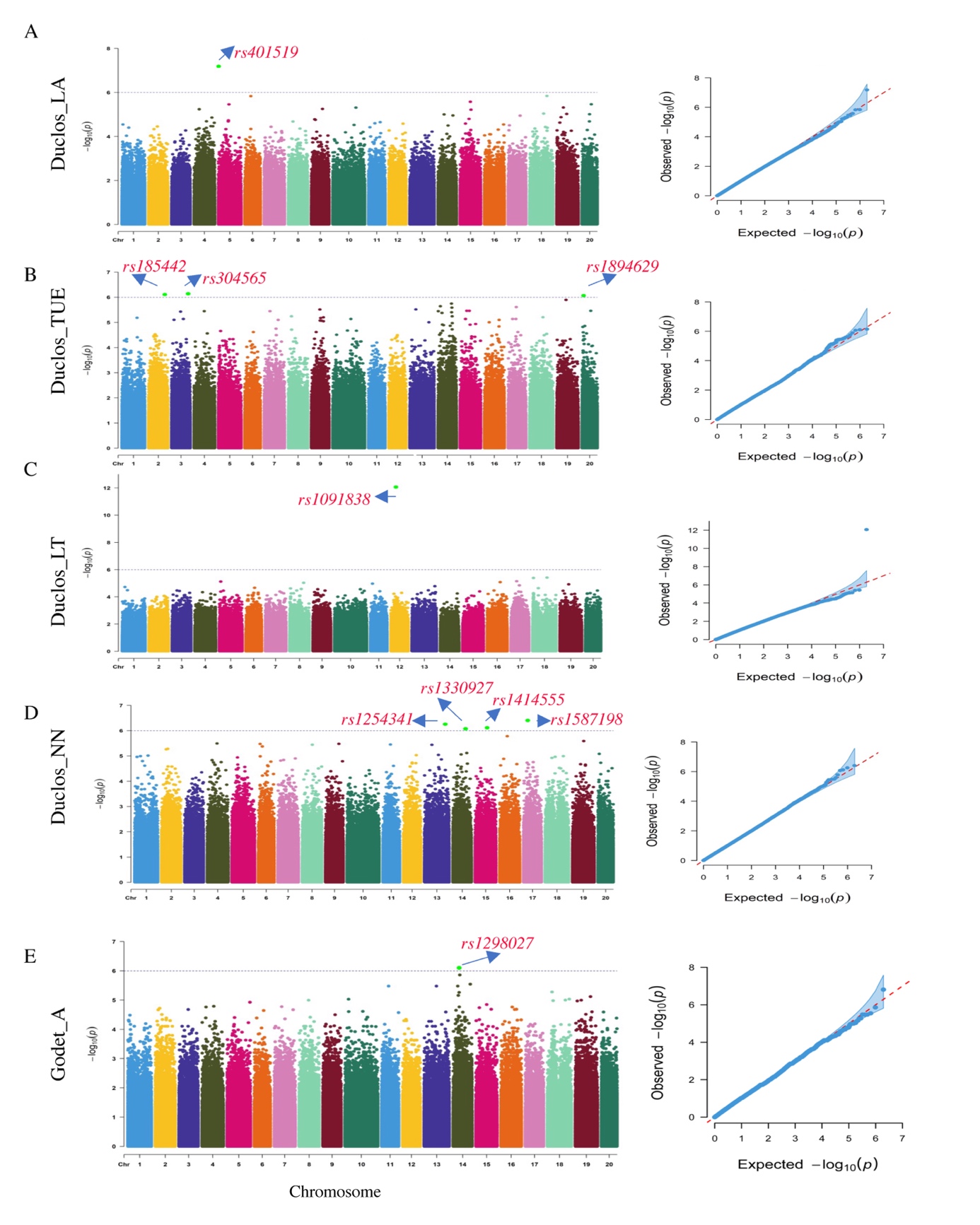

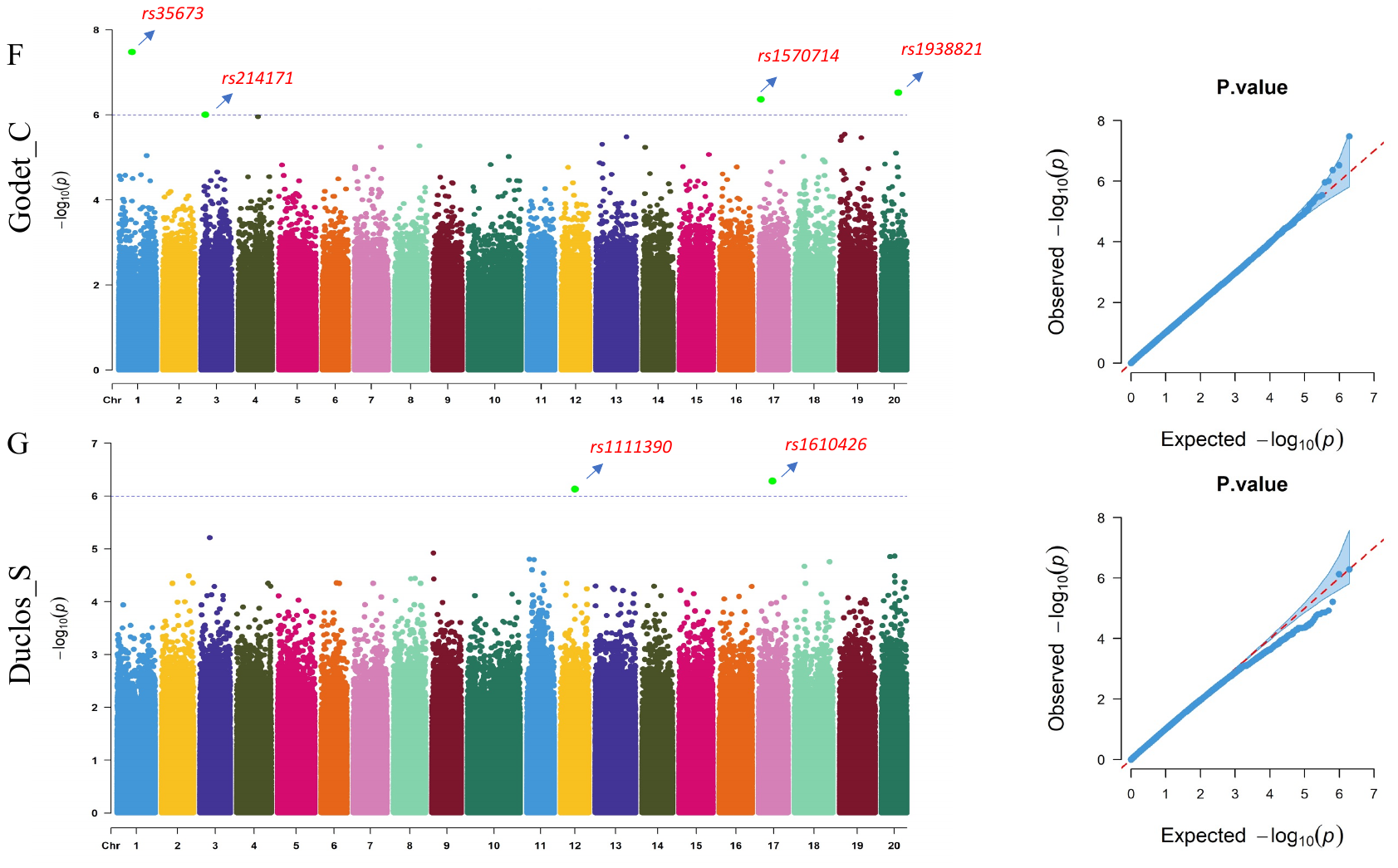

**Figure S3.** GWAS for traits under study. A) Manhattan plots for leaf area at Duclos with the peaks indicating significant GWAS signals and the dotted horizontal lines indicating the genome-wide significance threshold. While the right panel is a QQ-Plot associated with LA at Duclos, shows the -log_10_*P* of the expected vs. observed P values of each SNP (blue dots), B) Manhattan plots for transpiration use efficiency at Duclos, C) Manhattan plots for Leaf thickness at Duclos, D) Manhattan plots for Node number at Duclos E) Manhattan plots for net photosynthesis at Godet, F) Manhattan plots for competitor at Godet, G) Manhattan plots for stress-tolerator at Duclos. LA=leaf area, A=net photosynthesis, E= transpiration rate, TUE= transpiration use efficiency, NN= node number, LT= leaf thickness, C = competitor, S = stress-tolerator ecological strategies.
