## Supplementary material for "Identification and analysis of genomic regions influencing leaf morpho-physiological traits related to stress responses in *Dioscorea alata*": Figure S1

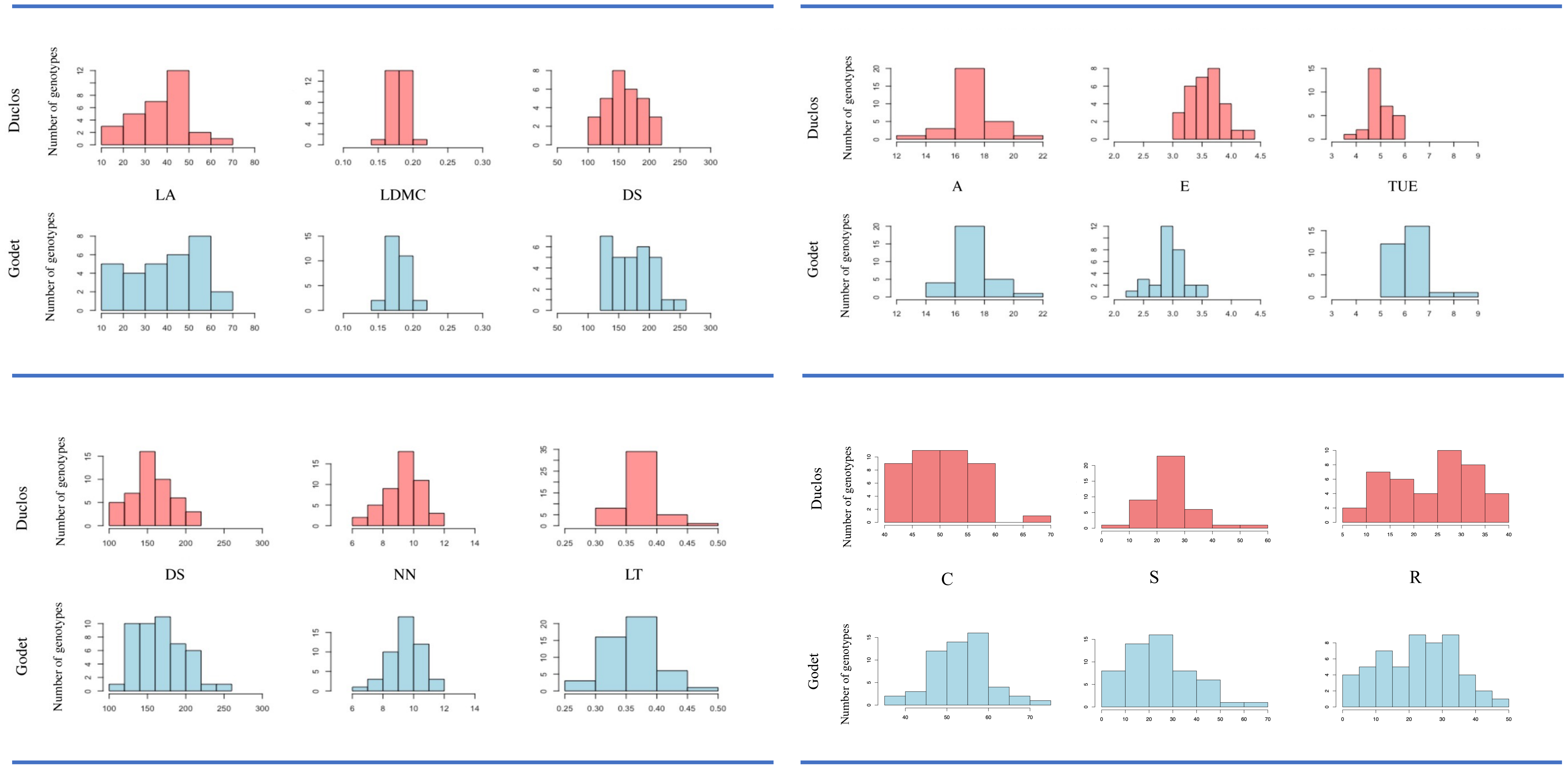


**Figure S1.** Distribution of each trait under study at two locations, Duclos and Godet. LDMC= leaf dry matter content, LA=leaf area, A=net photosynthesis, E= transpiration rate, TUE= transpiration use efficiency, IS= stomatal index, DS= stomatal density, NN= node number, LT= leaf thickness, C = competitor, S = stress-tolerator, R = ruderal
